# Nonmetric ANOVA: a generic framework for analysis of variance on dissimilarity measures

**DOI:** 10.1101/2021.11.19.469283

**Authors:** Alina Malyutina, Jing Tang, Ali Amiryousefi

## Abstract

Classic Analysis of Variance (ANOVA; cA) tests the explanatory power of a partitioning on a set of objects. Nonparametric ANOVA (npA) extends to a case where instead of the object values themselves, their mutual distances are available. While considerably widening the applicability of the cA, the npA does not provide a statistical framework for the cases where the mutual dissimilarity measurements between objects are nonmetric. Based on the central limit theorem (CLT), we introduce nonmetric ANOVA (nmA) as an extension of the cA and npA models where metric properties (identity, symmetry, and subadditivity) are relaxed. Our model allows any dissimilarity measures to be defined between objects where a distinctiveness of a specific partitioning imposed on those are of interest. This derivation accommodates an ANOVA-like framework of judgment, indicative of significant dispersion of the partitioned outputs in nonmetric space. We present a statistic which under the null hypothesis of no differences between the mean of the imposed partitioning, follows an exact *F*-distribution allowing to obtain the consequential *p*-value. Three biological examples are provided and the performance of our method in relation to the cA and npA is discussed.

**Significance Statement:** The Nonmetric Analysis of Variance (nmANOVA) conveys a framework that allows a compatible type of ANOVA for the cases where the proper metric measurements between objects are either lost, unknown or however inaccessible. While classic ANOVA is based on the measurements of the data from a base datum, the nmANOVA is formulated on the dissimilarity outputs (not necessarily metric) defined between all objects. As the main goal of ANOVA in providing a statistical test for assessing the significance of a considered partitioning on the data, the nmANOVA is yielding a paralleled scheme of inference with 1) accommodating the outcomes dissimilarities into *within* and *between* groups statistics, 2) assessing their respective divergence with a parametric distribution, and 3) providing a resultant *p*-value indicative of evidences fore rejecting the null hypothesis.

---

**C**lassic analysis of variance (ANOVA; cA) (1), stands as a renown generic statistical tool for testing mean similarity of different groups (2, 3). Since its introduction, cA has been developed to take into account various aspects of the experimental design and properties of observed variables from unbalanced sampling (4) to non-Gaussian residuals by Kruskal-Wallis type of nonparametric testing (5–7). A special case where the reference point to which the measurement of objects is not available but still one is able to collect all the pairwise distances between them, caught considerable attention by a series of work (8, 9). In this scenario, the emphasis has been put on the use of distances between responses rather than their actual values (9), as well as different measures for quantifying the distances (10), and a permutative method for forming a pseudo-*F* hypothesis testing statistic (hence nonparametric ANOVA; npA) based on the distances of the responses (11). In npA, the original sum of squared distances of responses from their means was replaced by the mean sum of square distances between responses. This was shown to lead into a variance decomposition that equates the case where the observation vectors themselves are considered (S3.2). While the quantification of most experimental variables could be measured with metric functions, there are cases for which the use of these functions are not more than a mere coercion. For example, a data transformation of non-metric outputs to Euclidean metric (12) or by incorporating a deduced metrics from a principal coordinates analysis on the dissimilarity matrix (11), while permits the npA model to operate, is leading to invalid judgments due to compressing and deforming the data beyond sufficiency. The failure to appropriately acknowledge the data inherency, is at the core of the misjudgment based upon the deficient data coercion. Network graphs rising in multiple biological studies for example (13–15), could exhibit the nonsymetrical measurements between the nodes, lack the edge between two, or even contain a self-looped nodes; all of which being cases where the metric properties are negated (S1). Each of these nonmetric scenarios should then be represented with a dissimilarity matrix between the nodes (rather than being metric-forced) and so demand a statistical framework that could incorporate this input as it is.

Here, based on the central limit theorem (CLT) and harnessing the natural relationship between statistical distributions (16), we generalize the cA method to a nonmetric case by introducing a nonmetric ANOVA (nmA). This development allows the direct integration of the nonmetric data input and unlike npA, provides a parametric *p*-value based on the theoretical distribution of an *F* -statistic that under null hypothesis of means similarity between partitioning classes follows the exact *F* -distribution. This enables the hypothesis testing for the experimental settings with nonmetric outputs. As we demonstrate with three diverse biological case studies, the nmA provides a statistical testing solution that allows the integration of the unmodulated dissimilarities input into a test statistic, downstream testing mechanics, and a resultant *p*-value.

## nmA tests differences of dissimilarities

In many practical cases the level of dissimilarities expressed between items does not follow the metric properties (17). Based on triangle equality, the simultaneous affirmation of all derived equations presented in 10 is necessary for a set of dissimilarities to be defined as metric (18). The negation of each to all of these equations (and hence invalidating the metric definitions) is extensively classified (19). For example, the term *semimetric* denotes scenarios where the triangle inequality fails to hold in general, while a *pseudometric* refers to cases where the forward condition of the identity property fails (20); the case where two distinct objects have a zero distance. Referring to the network analogy, this equates the cases with two distinct nodes with a complete connectivity such that the information in either of nodes is surely available in the other. *Metametric* on the other hand, is an opposite case where the distance between two identical items can be more than zero; a self-looped node that might loose the information received (21), and lastly, *quasimetric* refers to cases where the symmetry condition is distorted such that the distance from objects *A* to *B* is not necessarily the same as distance from *B* to *A* (22); such as inequality of the edges and hence unbalanced information flow between two nodes in a directed network. These violations of the metric properties in general provide a ground for assessing cases where the measurements do not exhibit metric properties (23). Next to the classic invention of nonmetric methods like Kruskal multidimensional rank test (24) and Mantel permutation tests (25), the prevalence of the nonmetric data has led to considerable advancement in other statistical domains as a proportionate response for the existing demand. For example, decision trees finding based on nonmetric data provides a robust tree optimization, and partial least square and generalized linear models, offer solutions for the regression fitting for nonmetric outputs (26–29). In alignment to these advancement and reference to the partitioning scenarios, here we extend the cA model to nmA. This extension allows for example, obtaining the *p*-value for each of the bifurcating nodes on the decision tree and assessing the multipartitism of a generic network. Also as a meaningful add to the nonmetric generalized linear models, nmA permits a similar hypothesis testing between objects in different partitions. As such, given the dissimilarity measurements, the resultant *p*-value of this method could also be used as a yardstick for finding the most distinct partitioning underlying a set of objects. (30).

## *F*_*nm*_ is a sufficient statistic for hypothesis testing

Consider an experiment where the independent variable imposes a partitioning on the responses and the task is to assess the significance of this partitioning. Similar to the nonparametric model (S3), the items in each partition are assumed to be exchangeable, but the dissimilarities between responses is now assumed to be any type of dissimilarities which are not necessarily outputs of a defined metric function.

Under the null hypothesis of similarity of all partition means, the *F*_*nm*_ as presented below is following the *F* - distribution with *g*^2^ − *g* and *g* − 1 degrees of freedom (S1);

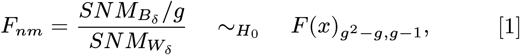

where 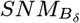 and 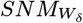 are the *sums of squared normalized means* of dissimilarity values *between* and *within* partitioning groups (S1).

The *F*_*nm*_ formulated in (2) is encapsulating the ratio of divergence of the mean of *between* to *within* groups. Under null hypothesis of no difference between the mean values of the partitioning groups, the sum of normalized means of dissimilarity values *between* each partition would be the same as their *within* counterparts (S2). To the degree of deflection from this ground truth, the observed *F*_*o*_ would be inflated and so the 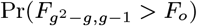 quantity would be the assigned *p*—value to indicate whether this deflection is of any significance with a designated *α* level threshold. Table 1 summarizes the essentials of this model with respect to the metric ones.

**Table 1.** Summary of different ANOVA models^*^.

| Method | ANOVA | Nonparametric | Nonmetric |
| --- | --- | --- | --- |
| Specification | Responses | Distances | Dissimilarities |
| Input | $\mathbf{y}$ | $D_{N \times N}$ | $\Delta_{N \times N}$ |
| Test Statistic | $\frac{SS_B/g-1}{SS_W/N-g}$ | $\frac{SS_{B_d}/g-1}{SS_{W_d}/N-g}$ | $\frac{SNM_{B_\delta}/g(g-1)}{SNM_{W_\delta}/g-1}$ |
| $P$-value | $P(F_{g-1, N-g} > F_o)$ | $\frac{\sum_{\pi} 1_{[pF_{np}^{\pi} \geq pF_o]}(\pi)}{\sum_{\pi} 1}$ | $P(F_{g^2-g, g-1} > F_o)$ |

\*The high level similarity of nmA to cA is its closed form distribution of the Test Statistic leading to the respective $P$ -value, while the matrix structure of the Input is common between npA and nmA, albeit former based on the Distances( $D$ ), and Dissimilarities ( $\Delta$ ) the latter (S1-S3).

## General properties of nmA

### nmA is sensitive to groups differences

To assess nmA performance in different situations, we conduct series of simulations from Gaussian and uniform distributions which cover various scenarios via adjusting distribution parameters. While literally any other parametric distribution could be harnessed in here, this choice of models was encouraged as at the constant variance level, the Gaussian distribution is known to have the minimum negentropy (31, 32); hence the most chaotic sequences of values are expected. The uniform distribution on the other hand is chosen as it is usually taken as the state of uninformativeness in the Bayesian scenarios (33).

Firstly, we are interested in the performance of the nmA when the null hypothesis is true. Under this condition for any parametric test, the *p*-values should follow the *U* (0, 1) distribution (34). In our case, using the same sampling distribution for generation of the dissimilarities with given partition imposed on them led to the consistent conclusion (Figure 1A).

**Fig. 1.**
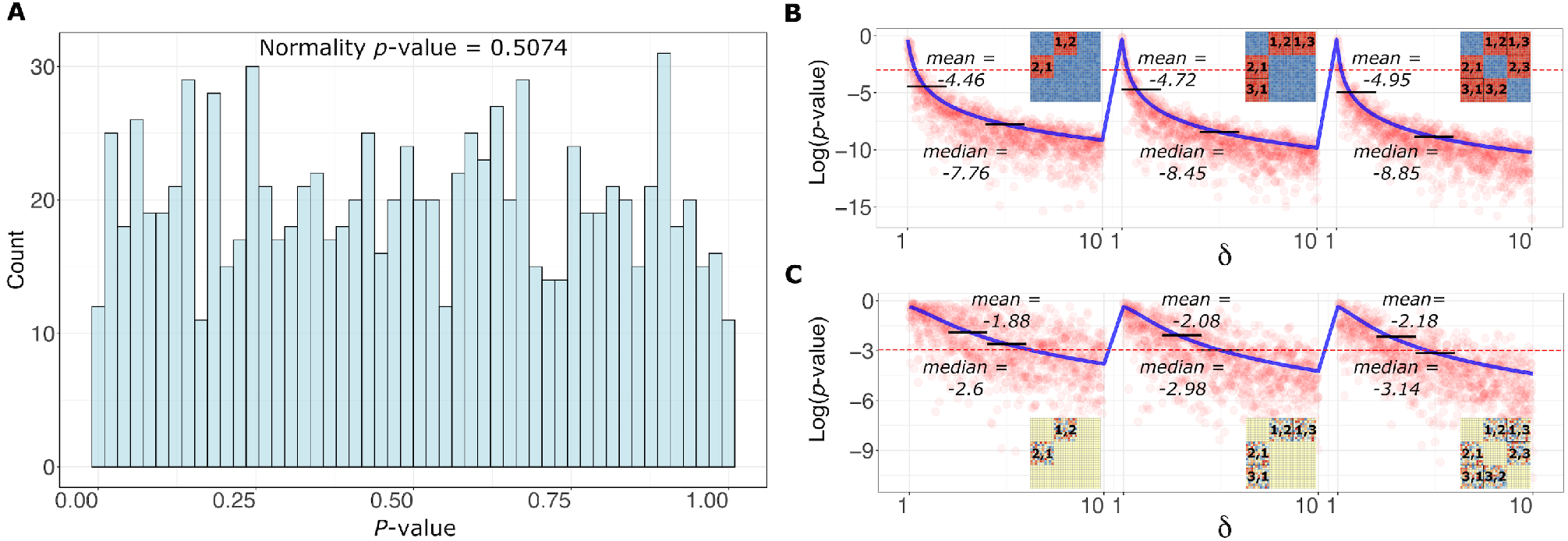
The consistency of the *F*_*nm*_ under different scenarios. **A)** *Uniform distribution of the p-values*. Uniform distribution of 1,000 *p*-values for three equal-sized groups of 10 when the whole dissimilarity matrix is coming from *U* (0, 1) distribution. The results of two-sided Kolmogorov-Smirnov test are highlighted on top. The same behavior was observed with simulation from normal distribution with 4 groups of 10 each and 1,000 replication (*p*-value = 0.635). **B)** *The effect of progressively changing the mean of the between group matrices distribution on the p-values*. We use the previous group dissimilarity matrix as our starting point and continuously change one of the uniform distribution parameters via adjusting *ψ* factor (as an incremental adjustment of the *a* and *b* in *U* (*a, b*)). With every step *i* from [1, 10], we change the *between* group matrix distributions to be *U* (0 + *ψ*_*i*_, 1 + *ψ*_*i*_) and obtain the corresponding *p*-value. At the same time, we track the effect of the proportion of *between* group data being affected via *ψ* parameter. The three scenarios are visualized (from left to right): when only group 1 and group 2 pairwise *between* group matrices are adjusted; when group 1 and group 2 but also group 1 and group 3 dissimilarity matrices are simultaneously adjusted; when all the 6 possible *between* group matrices are simultaneously adjusted. The red dots represents each *p*-value with corresponding *ψ* and the mean values of them are connected with the solid blue lines. The red dashed line corresponds to the *p*-value equal to 0.05. **C)** *The effect of progressively changing the standard deviation of the between group matrices distribution on the p-values*. While in previous case the difference in *between* group distributions was triggered via changing the mean of uniform distribution, here we fix the mean but change standard deviation via adjusting the *between* group distribution according to *U* (0 *− ψ*_*i*_, 1 + *ψ*_*i*_). The adjustment is performed for the same three scenarios as previously. As indicated with the lower values (of the mean and medians) and the higher gradient of descent (solid blue lines), nmA is more sensitive with the dislocation of the mean as the first central moment of the values than the dispersal standard deviation parameter.

Secondly, moving toward the alternative hypothesis, we investigated the sensitivity of the model with respect to the marked changes in the means of partitioning groups as a measure of deviation from the null hypothesis. The incremental shifting of the values of the *between* groups with respect to the *within* groups has led to the expected inflation of the *F*_*o*_ statistics in nmA model (Figure 1B). Despite sensitivity of nmA to the location parameter, the scale changes were not leading to the same significant effect as increasing the standard deviation of *between* to *within* groups does not lead to the marked changes in the nmA judgment denoting the robustness of the method with respect to scale dimension of the inputs (Figure 1C).

In contrary to its conventional use, nmA can assist in selecting the most similar sub clustering of the objects aggregated in a dissimilarity matrix. This application of the method was assessed by tailoring the apparent differentiation between the groups and allowing the model to retrieve that (Figure 2A). To do that, we first simulate a dissimilarity matrix which represents 4 groups of different sizes - 13, 21, 34, 55. Then we split the matrix into 3 groups via joining the first two clusters, or 5 groups via splitting the last partitioning into 2 groups of sizes 21 and 34. We fix *within* and *between* group dissimilarities distributions to *N* (0, 1) and *N* (5, 1) respectively. It is important to note that we select the cluster sizes on purpose, as they represent a sub-sequence from a Fibonacci sequence allowing minimum inclusion of the *between* elements to *within* when joining or splitting the clusters. Fixing the umber of clusters to 3 as described, we obtain clusters of 34, 34 and 55 sizes. This will not only provide relatively similar sizes for the clusters but also guarantee the smallest influence on the *between* original cluster 1 and cluster 2 dissimilarity values. More precisely, we get 2*13 × 21 *between* cluster 1 and cluster 2 dissimilarity values coming from *N* (5, 1) distribution mixed with 13^2^ + 21^2^ dissimilarity values from *N* (0, 1). Thus, organising clusters this way lead to the smallest possible influence of higher-valued (*N* (5, 1)) dissimilarities to the newly-obtained *within* group matrix. Similarly, via splitting the cluster of size 55 into 2 clusters of 21 and 34, we provide the minimum values from *N* (0, 1) to appear in *between* group dissimilarities. However, the *p*- values that correspond to the 5 group partitioning are very close to the initial partitioning *p*-values, but their difference starts to grow with the increase of elements rearrangement (Figure 2A). The *p*-value closeness in the beginning can be explained via subtle changes in *between* group matrices with the strategy we selected for partitioning. Overall, progressive cluster rearrangements were reflective of growing *p*-values to the level that no more differences between imposing groups were existing. The resulting *p*-values have indicated that no partition can achieve as low *p*-values as those that we got with the initial most differentiated clustering (Figure 2A).

**Fig. 2.**
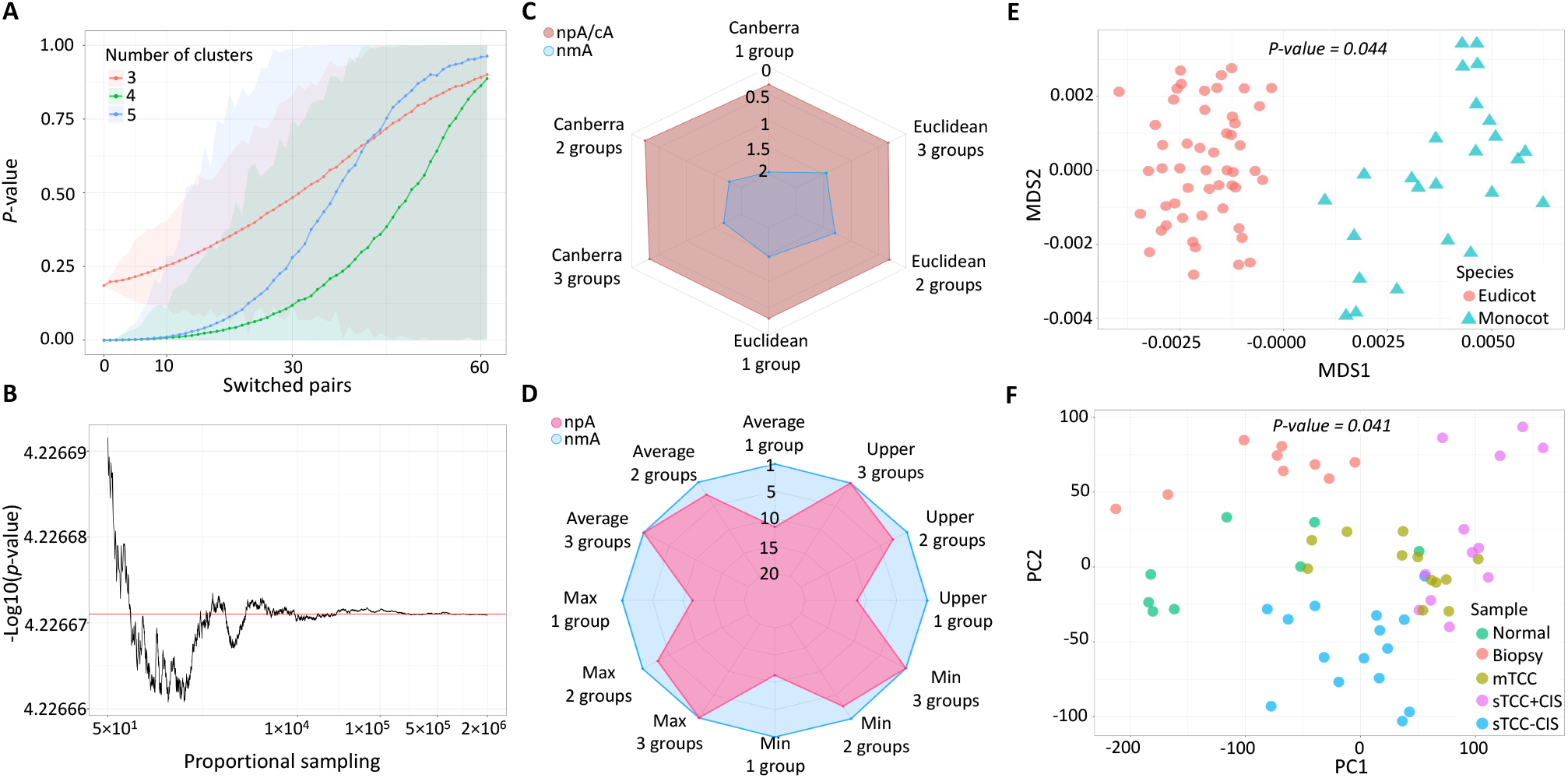
nmA properties and applications. **A)** *Distribution of p-values to define optimal clustering*. The original dissimilarity matrix clustering of size 4 has been rearranged to size 3 and 5 to demonstrate that nmA’s *p*-value can serve as an indicator for selecting the optimal clustering of the data. Additionally, there has been up to 60 between cluster pairs switches to make the fixed clustering shuffled. The dotted lines represent the average *p*-values over 1,000 scenarios of switched pairs. The shaded area spans between the minimum and maximum of the *p*-values obtained for each case. **B)** *The convergence of p-values with asymptotic sampling*. The *p*-values converge to their mean value (highlighted in red) after approximately 10^4^ proportional samplings performed. **C)** *nmA is more conservative with metric inputs*. Four vectors with 100 elements were created using *N* (10, 1) for cA and either Euclidean or Canberra distance was applied to them to obtain 400*×*400 distance matrix for npA. To make nmA applicable, the matrix was transformed via placing the elements of its upper triangular part randomly to the upper and lower triangular parts of a new matrix and taking into account which grouping the elements belong to. The values that has not been imputed in this manner are treated as missing. We further add a small noise from *N* (0, 5 *×* 10^−4^) to all the values of the matrix. After that, the mean of either one, two, or three vectors was progressively increased from 0 to 2. The figure shows the space where a method considers the current data partitioning significant. While cA and npA congruently detect differences at a smaller mean differences, the nmA is exhibiting a more conservative trend. **D)** *nmA outperforms cA and npA in nonmetric cases*. The 400*×*400 matrix with *within* and *between* group matrices being from *N* (30, 1) and *N* (10, 1) distribution is used for nmA and its the coerced symmetric version for npA. The symmetry was achieved via either taking average, minimum, or maximum of the two symmetric counterpart values of the matrix or utilizing the upper triangular part of the created matrix as the input for npA. Similarly to previous case, we start to increase the mean of either one vs two, two vs three or three vs four *between* group matrices while keeping the *within* group matrices the same. This time the result indicates the sharper detection of the differences between the means for the nmA compared with npA. **E)** *MDS bipartitioning of the BLAST data*. The MD applied on the distance matrix obtained for every species pair as an average of their two nonsymmetric dissimilarities. The reciprocal BLAST MDS clearly separated the data into monocots and eudicots. nmA confirms this grouping with a *p*-value = 0.044. **F)** *PCA-based grouping of the bladder gene expression data*. Some of the sample groups are separated from others while some are rather mixed. nmA provides a significant *p*-value = 0.041 as it is sensitive even to one *between* group difference in the data.

**Fig. 3.**
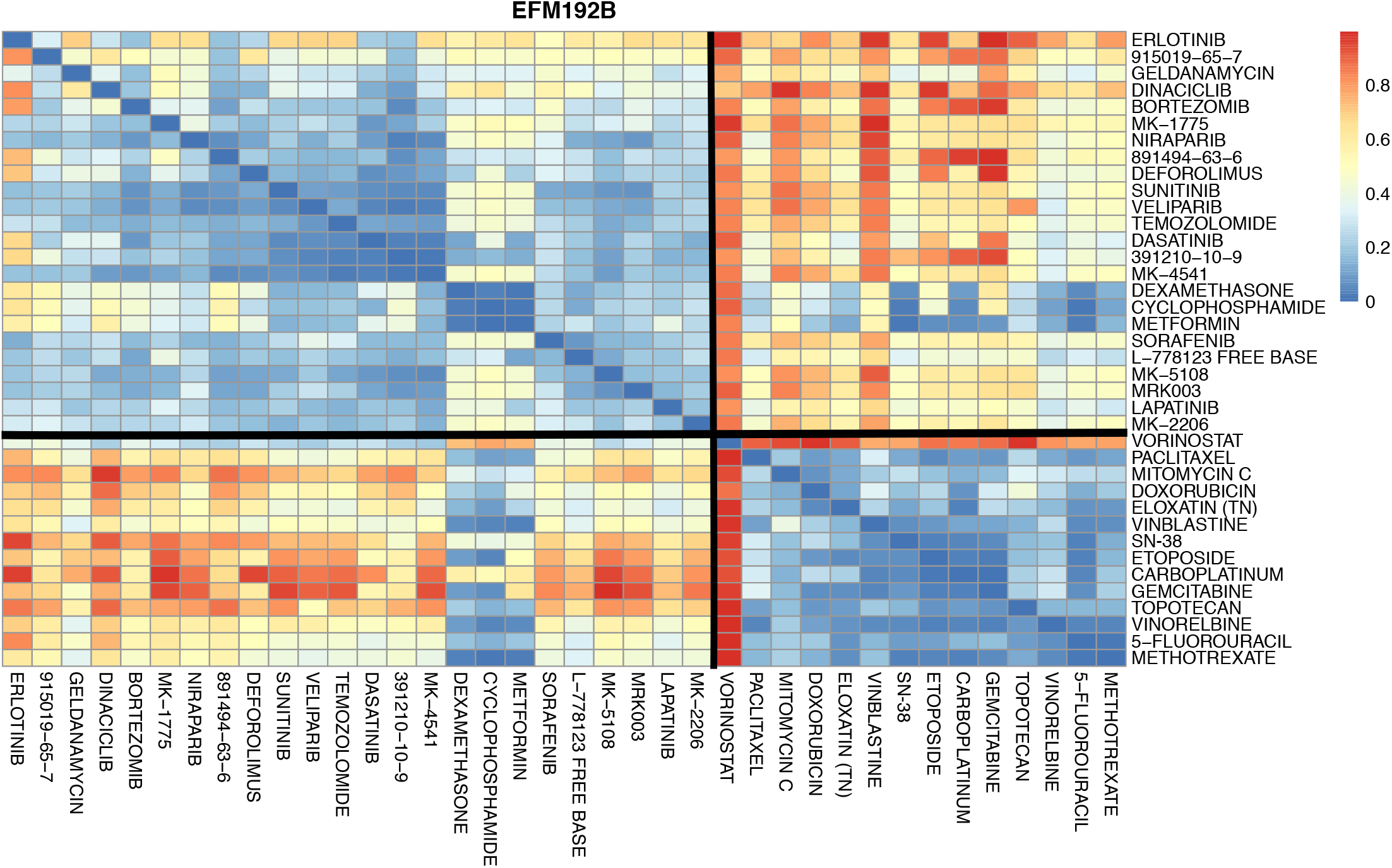
Significant bipartitioning of CSS-based dissimilarity matrix for EFM192B cell line. The dissimilarity matrix is achieved by first determining the quasimetric 38 *×* 38 CSS-based matrix with rows and columns corresponding to the drugs. At the same time, we fill the diagonal values with RI scores, as they bring additional information about the drugs’ sensitivity. Then obtaining two directional correlation-based matrices; we subtract the absolute values of a correlation matrix applied to either CSS-based matrix or to its transpose version, from 1. After that, we create a joint matrix via filling upper and lower triangular parts of the new matrix with the upper triangles of the obtained symmetric matrices. Thus, we take into account correlation-based drug similarities as nonsymmetric. We also fix diagonal values of the matrix to zero given that the row and column drug is the same. The obtained EFM192B specific bipartitioning, which is marked by the two black lines on the dissimilarity matrix based heatmap, resulted in *p*-value = 0.034.

**Fig. 4.**
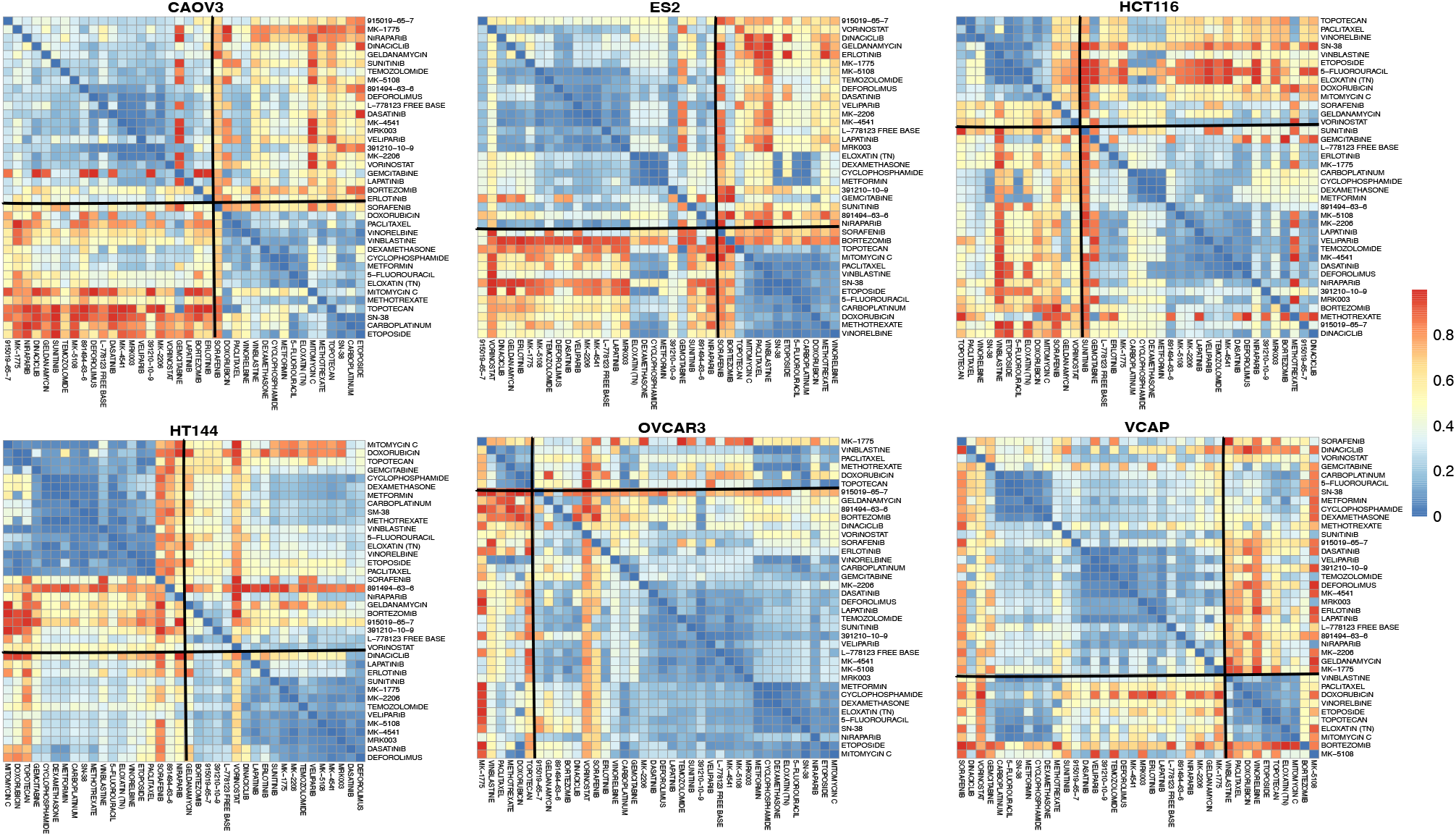
Significant bipartitioning of CSS-based dissimilarity matrices for 6 cell lines. The marked bipartitioning of the cell line specific dissimilarity matrices which are represented via heatmaps, has been marked as significant by nmA.

**Fig. 5.**
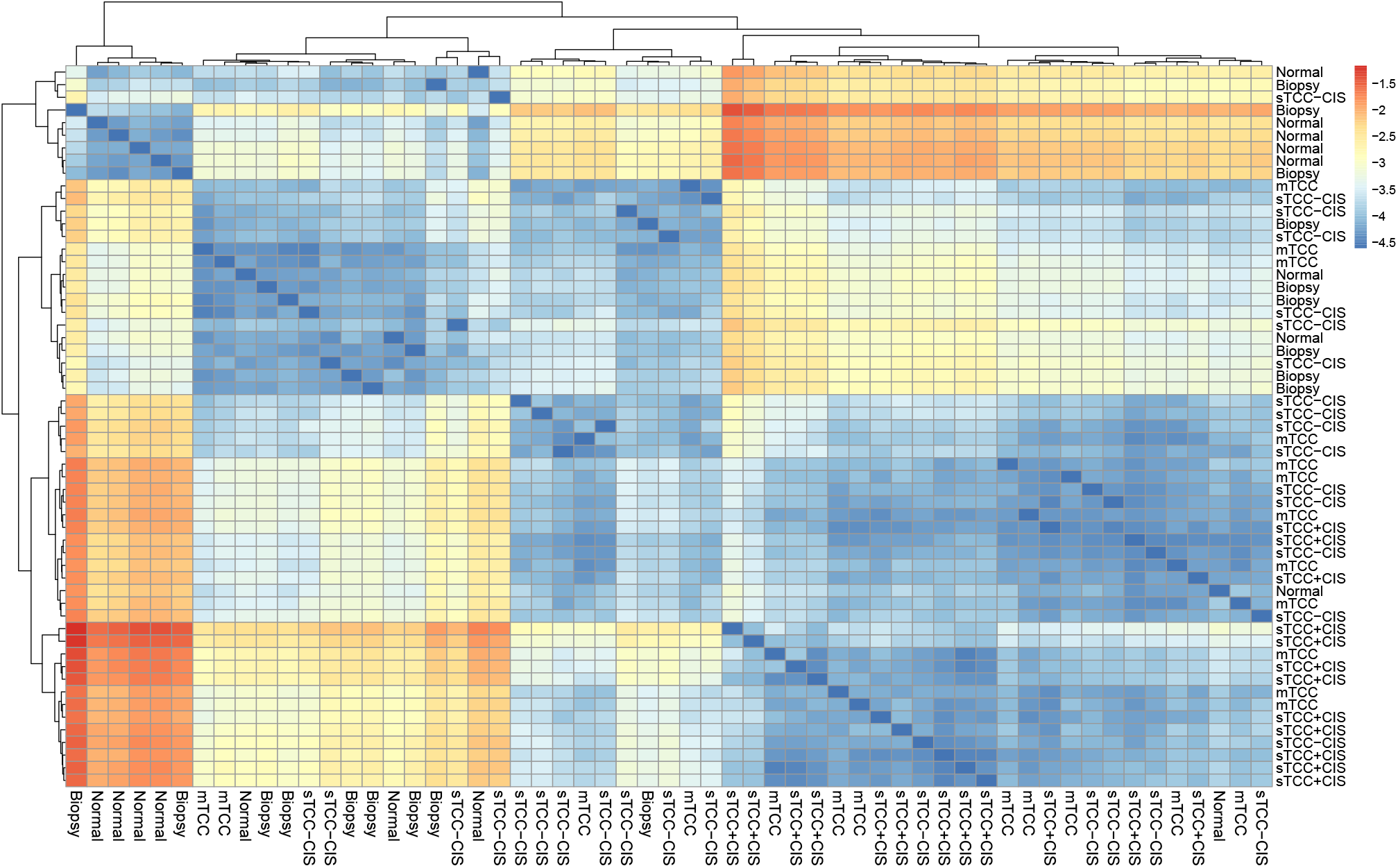
KL divergence-based dissimilarity matrix for the bladder gene expression data. In order to obtain a dissimilarity matrix for the data, we apply KL divergence scoring. We first transform sample specific gene expression distributions into probability distributions and then obtain KL divergence values for each distribution pair. Since KL divergence calculation is dependent on the order of the two distributions, it is non symmetric. We visualize the obtained dissimilarity matrix via a heatmap of its logarithmic values. We also apply unsupervised hierarchical clustering to show relation between the samples. We find out that dissimilarity matrix separates 3 carcinoma groups (mTCC, sTCC-CIS and sTCC+CIS) and Normal or Biopsy samples. These separations have been marked as significant by nmA with *p*-values listed in Table 5.

### Asymptotic sampling improves accuracy

To assess the convergence of *p*-values with increasing number of the proportional samplings (S1), we perform simulations where we collect a sequence of *p*-values which correspond to the same dissimilarity matrix but with a varying number of proportional samplings. Every *p*-value in a sequence is obtained with one more proportional sampling added to the one obtained for a previous *p*-value. For these simulations we select 3 groups of sizes 10, 20 and 30 and sample dissimilarity matrix from *U* (0, 1) for *within* group matrices and *U* (2, 4) for *between* group matrices. Thus, we introduce a difference between the groups via increasing mean and standard deviation, and expect *p*-values to reflect it. To check the asymptotic sampling behaviour, we apply nmA to the matrix and increase proportional sampling number from 1 to 2 × 10^6^. As Figure 2B shows, *p*-values capture the significant difference between the groups and converge to their mean value. This example demonstrates that one can improve the accuracy of a *p*-value via increasing the number of proportional samplings while even a small number of proportional samplings could be conductive of a significant difference.

### nmA outperforms cA and npA in nonmetric cases

We also consider the comparative performance of cA, npA and nmA. We note that while npA reduce to cA with the Euclidean inputs (9), the relation of the nmA to either cA or npA is not canonical. So any type application of the nmA method on the metric data would not be more than a coercion and hence the results would not be comparable nor conductive of any judgment. Here however, we were interested in examining some of these main data modulation that at least structurally formats the date to be a transferable input across different models. The results showed the almost identical sensitivity of the npA and cA models under two different metric measures while the application of nmA on the modulated data led to a more conservative threshold for the rejection of the null hypothesise (Figure 2C). The expected congruency and high sensitivity of the cA and npA models are mainly due to the correct application of these models on the very data that are within the space of legit inputs of these models. However, nmA being abused on the metric data, our result underlies the rather more conservative sensitivity with regard to the null hypothesis, such that significant judgment for the rejection of it is always supported with the cA and npA while not necessarily the other way around (Figure 2C).

Tailoring the inputs perfectly for the nmA model and this time coercing the data into metric, fit for the npA model, we observed an almost inverse phenomenon (Figure 2D). With the nonmetric data, nmA is more sensitive than the npA with respect to rejection of the null hypothesis. We believe the same reason of inadequate compression of the inputs to form a proper input candidate of the npA model is again the main cause of this discrepancy. In effect, this is confirming the merit of each models in the defined sphere of their underlying assumptions with regards to the input data.

## nmA provides solutions for real datasets

### Phylogenetic bipartition assessment with BLAST bit scores

For our first example, we consider a set of 78 angiosperms - flowering plants that bear their seeds in fruits (35). The data is split into two groups: 26 monocots (one seed leaf plants) and 52 eudicots (two seed leaf plants) (Table S3), leading to a total of (78^2^) 6084 possible pairings. For each pair, we collect basic local alignment search tool (BLAST) output bit scores of the fundemantal chloroplast conserved ribosomal protein S8, *rps8* gene. Since BLAST bit score is a semimetric measure (36), the cA and npA are not applicable, as neither a base value for a single gene is available (cA requisite) nor the bit scores for angiosperms represent a metric (npA requisite). A higher bit score is a marker of genetic homogeneity between the species, therefore inverted bit scores could be used as dissimilarity measurement of their phylogeny. Aggregating these values for all the possible pairs of the species led to the nonmetric matrix of dissimilarities (Δ _78×78_; SD1). Considering the partitioning of species into monocots and eudicots imposed on the above matrix, we quantify the significance of this bipartition via applying nmA. Our method indicates that the difference between the two species groups is significant with a *p*-value of 0.044 (*F*_*o*_ = 257.87), which is in accordance with prior findings in plant physiology (37, 38). Indeed, the separation is noticeable on Figure 2E, where we apply multidimensional scaling (MDS) to the data matrix (39). This result suggests the use of nmA for the phylogenetic bifuricating structure assessment that is often based on nonmetric similarity matrices. The higher order indices of nucleotide sequence similarities (e.g. bit score, E-value (40)) are obtained from an underlying nonmetric nucleotide substitution matrices. In fact, all the nucleotide substitution models that allows for nonsymetric nucleotide substitution rates (F81, HKY85, TN93, and GTR) will essentially result in nonmetric similarity measure between sequences (41). As the metric conditions can be also negated in the case of the codon or amino acids substitution models (such as BLOSUM or PAM), the nmA use can be extended for downstream assessment of partitionings constructed on these models outputs as well (42–45).

### Drug combination sensitivity scoring for cell lines

Combination sensitivity scores (CSS) have been introduced to provide information about drug combination efficacy applied to cancer cells (46). After drugs sensitivity estimation, it is usually of interest to see whether particular drug groups lead to similar sensitivity patterns. The directional nature of the CSS scores as to which drug combination is a baseline, renders the CSS quasimetric. Consequently, nmA can be applied to any partitioning of drugs to evaluate its significance. In this example, we apply nmA to assess significance of drug clusters obtained from drug combination experiments. We download relative inhibition (RI) scores (which are sensitivity scores in single drug experiments) and CSS from DrugComb portal (47). We filter data to come from O’Neil drug combination study as it contains the most complete dataset (48). The data provides RI and CSS for 38 unique drugs and 39 cancer cell lines (SD2). For each cell line, we obtain dissimilarity matrices for the 38 drugs, based on the correlations of their CSS_1_ or CSS_2_ values (Figure S3). Afterwards, we apply hierarchical clustering on the obtained matrices and fix the number of clusters to 2 aiming to find the most distinctive drug partitioning. To avoid cases when clustering is governed by a small number of drugs (e.g. due to drugs exceptional sensitivity patterns), we apply nmA to dissimilarity matrices that have at least three drugs in each partition. As a result, we identify 7 cell lines out of 39 for which the obtained partitioning is significant (Table 2, Figure S4). For example, the 38 drugs are clearly separated into two clusters for EFM192B cell line (Figure S3), which can be explained by their mechanisms of actions as the majority of the first group compounds are targeted drugs, meanwhile the second group mostly consists of conventional chemotherapy drugs; antineoplastic agents, mitotic and topoisomerase inhibitors (Table S4). The same trend can be seen for the remaining 6 cell lines (Figure S4), which supports the use of nmA as a tool for assessing the significance of the proposed clustering.

**Table 2.**
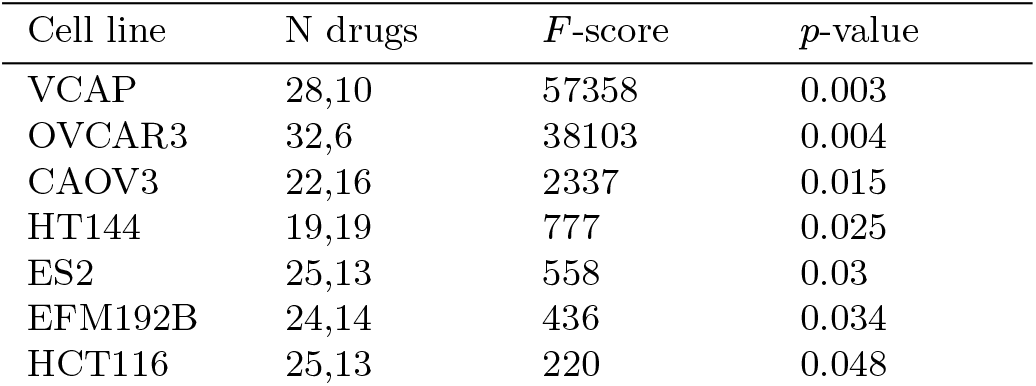
Nonmetric ANOVA model specifications that shows significance (*p <* 0.05) for the CSS data.

**Table 3.** Set of 78 angiosperms partitioned into 52 eudicots and 26 monocots that was used to form a semimetric matrix of dissimilarities based on the inverse values of the bit scores of *rps8* chloroplast gene for all the possible pairwise comparisons.

| Eudicots | Monocots |
| --- | --- |
| *Aethionema grandiflorum* | *Acidosasa purpurea* |
| *Ageratina adenophora* | *Acorus calamus* |
| *Anthriscus cerefolium* | *Agrostis stolonifera* |
| *Arabidopsis thaliana* | *Anomochloa marantoidea* |
| *Arabis hirsuta* | *Bambusa emeiensis* |
| *Arbutus unedo* | *Brachypodium distachyon* |
| *Atropa belladonna* | *Ferocalamus rimosivaginus* |
| *Barbarea verna* | *Festuca arundinacea* |
| *Betula pendula* | *Hordeum vulgare* |
| *Brassica napus* | *Indocalamus longiauritus* |
| *Capsella bursapastoris* | *Leersia tisserantii* |
| *Carica papaya* | *Lemna minor* |
| *Castaneamollissima* | *Lolium perenne* |
| *Citrus sinensis* | *Oryza meridionalis* |
| *Coffea arabica* | *Panicum virgatum* |
| *Crucihimalaya wallichii* | *Phalaenopsis aphrodite* |
| *Cucumis melo* | *Phyllostachys nigra* |
| *Daucus carota* | *Rhynchoryza subulata* |
| *Eleutherococcus senticosus* | *Saccharum hybrid* |
| *Eucalyptus grandis* | *Sorghum bicolor* |
| *Fagopyrum esculentum* | *Spirodela polyrhiza* |
| *Fragaria vesca* | *Triticum aestivum* |
| *Glycine max* | *Typha latifolia* |
| *Gossypium raimondii* | *Wolffia australiana* |
| *Guizotia abyssinica* | *Wolffiella lingulata* |
| *Helianthus annuus* | *Zea mays* |
| *Hevea brasiliensis* |  |
| *Ipomoea purpurea* |  |
| *Jacobaea vulgaris* |  |
| *Lactuca sativa* |  |
| *Lepidium virginicum* |  |
| *Lobularia maritima* |  |
| *Lotus japonicus* |  |
| *Manihot esculenta* |  |
| *Medicago truncatula* |  |
| *Millettia pinnata* |  |
| *Morus indica* |  |
| *Nasturtium officinale* |  |
| *Nelumbo nucifera* |  |
| *Nicotiana tomentosiformis* |  |
| *Oenothera elata* |  |
| *Olea europaea* |  |
| *Olimarabidopsis pumila* |  |
| *Panax ginseng* |  |
| *Parthenium argentatum* |  |
| *Populus trichocarpa* |  |
| *Prunus persica* |  |
| *Pyrus pyrifolia* |  |
| *Silene noctiflora* |  |
| *Solanum lycopersicum* |  |
| *Spinacia oleracea* |  |
| *Theobroma cacao* |  |

**Table 4.**
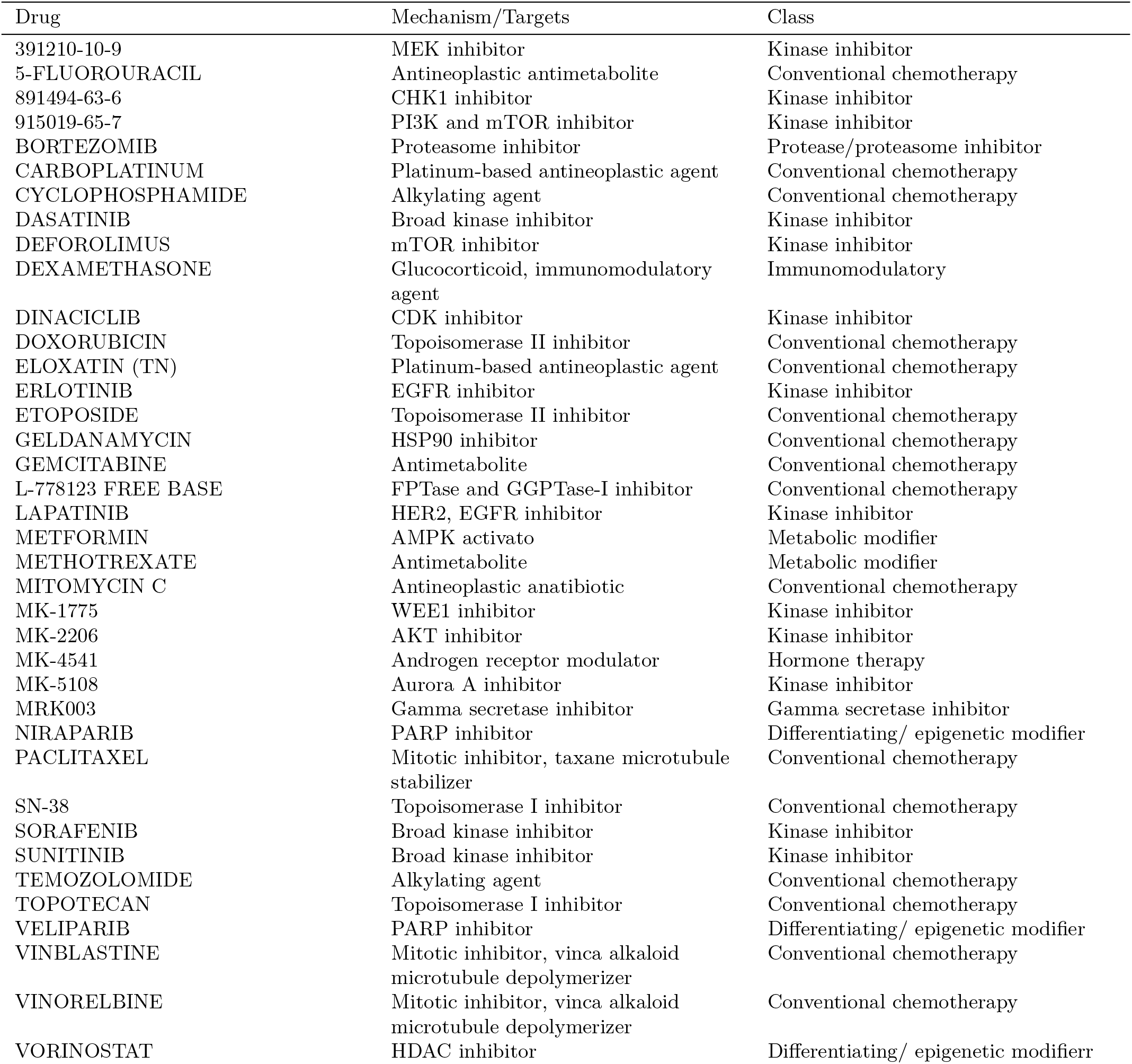
Mechanism of action/target and class information for the drugs used in CSS example.

**Table 5.**
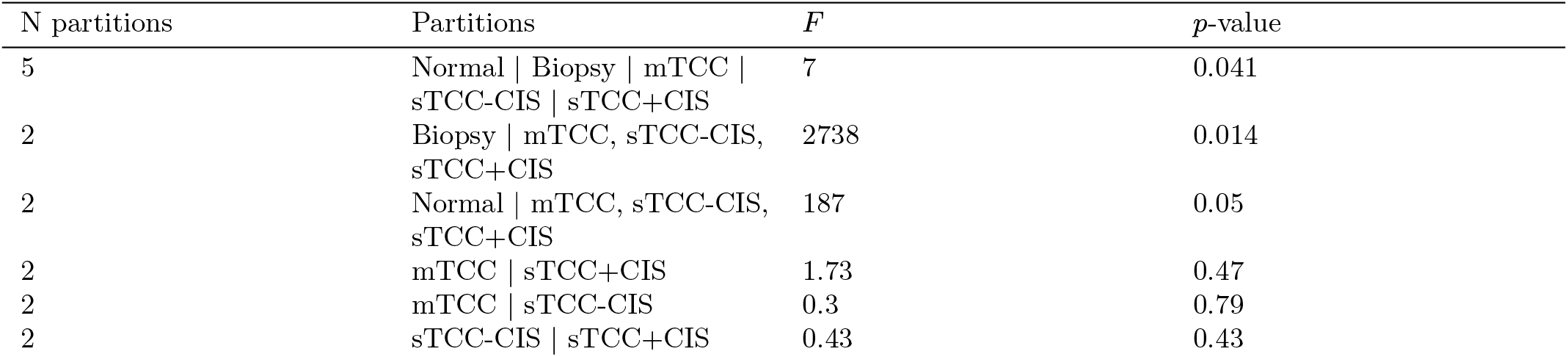
Nonmetric ANOVA model specifications for the bladder gene expression data.

### Kullback-Leibler divergence for gene expression data

For our next example we consider bladder cancer gene expression data obtained for 57 patient samples that were separated into 5 groups: 12 samples representing superficial transitional cell carcinoma with surrounding carcinoma *in situ* (sTCC+CIS) and 16 samples without surrounding carcinoma (sTCC-CIS), 12 samples within muscle invasive carcinomas (mTCC), 8 samples of normal bladder cells from healthy patients (Normal), and 9 histologically normal samples (Biopsy) close to the carcinoma *in situ* location (49, 50). We aim to apply nmA to check if the difference *between* the sample groups is supported with the gene expression-based dissimilarity matrix. To capture the full distributional level of proximity between the gene expressions, we used the Kullback-Leibler (KL) divergence (51) rooted in information theory. This choice is preferred over commonly known point-based methods of differential gene expression such as *t*-test as its providing a more comprehensive level of information regarding the dispersion of the expressions. We first create a dissimilarity matrix via applying KL divergence scoring on the bladder cancer gene expression data (N = 22,283). We apply unsupervised hierarchical clustering to the dissimilarity matrix (Figure S5) and principal component analysis (PCA) to the raw gene expression counts (Figure 2F) to check if KL-based clustering is in accordance with PCA grouping. It can be seen that KL-based dissimilarity matrix separates 3 carcinoma groups, namely mTCC, sTCC-CIS and sTCC+CIS, and Normal or Biopsy groups even better than PCA. Therefore, nmA distinguishes both Normal and Biopsy group as significant separation from 3 carcinoma groups (Table S5). At the same time, neither any bipartitions within 3 carcinoma groups nor Normal versus Biopsy partitions are found to be significant (Table S5). The initial partitioning of the bladder gene expression data into 5 groups is significant with *p*-value of 0.041. Similarly to cA, nmA test provides a significant *p*-value if at least one distinct bipartition is present in the data.

## Discussion

We introduced a generic type of nonmetric ANOVA method that is based on the dissimilarities between a set of objects. These dissimilarities are not needed to be defined with a metric function nor produced systematically. As long as a numeric value representing the closeness of a set of items subject to different partitions are given, the nmA could be applied for statistical quantification of the significance of the underlying partitioning. In cases where the clear partitioning of the data is not obvious or given *a priori*, the permutative use and search for the lowest *p*-value could lead to the most distinctive possible partitioning of the data such as the use case presented as the drug CSS example. In general, this method could be harnessed for quantification of any sort of clustering of a set of objects into different classes as explored in the first example, where it shows the possibility of the nmA for assessing the meaningfulness of a derived bifurcating structure on a phylogenetic tree (52). The other use could be on the network graphs where a specific multipartitism with an underlying dissimilarity matrix is given. As pointed in the last example, the nmA can provide a statistical testing for the patient stratification with reference to their gene expressions proximity. The same method of quantification could be applied for labeling the level of distinctiveness of cells in the UMAP or t-SNE plots based on the single cell differential RNA expressions measured with nonmetric methods such as KL divergence (53, 54).

The performance of the nmA to exiting methods including cA and npA also needs to be properly acknowledged. While the cA is solely constructed on the response values and supported by parametric assumptions such as normality of the residuals (7), the npA is formulated on the distances between the responses, free from parametric assumption but still confined to the metric conditions. Similar to npA, nmA is constructed on the dissimilarity measures between responses and devoid of any metric or parametric assumptions. Based on the CLT, the closed distribution of the test statistic for nmA is obtained, allowing a parametric derivation of the *p*-value. While proportional sampling could provide a more robust outputs for the nmA, we note that the comparative analysis across the discussed models is still primarily biased as lack of a totally objective method in transforming the data to fit perfectly to each model (Table 1). While these transformations are creeping the obtained results and hence, rendering the comparative task ineffective, the retrieval of the responses themselves (as needed for cA) is impossible even with the availability of the dissimilarities or distances between objects (Figure 2D, 1).

We also note the limitation of the nmA in face of growing number of partitions in relation to the number of observations. This is leading to quadratic explosion of the *between* samples values with regard to *within*. While the degree of the freedom division in *F*_*nm*_ formulation is accommodating for this, the different rate of convergence of the underlying means applied in the CLT is creeping the results toward a non-centrality of the *within* mean square error leading to a more false negative judgments toward rejecting the null hypothesis (S1). Similar phenomenon could be observed in the case of unbalanced partition sizes where the results would be primarily affected by the biggest partition. In such cases a weighted sampling can be a solution, however in extreme unbalanced scenarios this may not be tractable. Furthermore, the curse of dimensionality aroused in multivariate ANOVA and the unbalanced sampling are also challenging scenarios for cA that need to be properly addressed specifically in each experiment (55, 56) and likely so, nmA could be both developed and adopted in similar scenarios. Taken together,we provide nmA as an attractive method for a variety of scientific questions which involve the assessment of a given partitioning from nonmetric dissimilarity measurements.

## Data availability

The R code for implementing the cA, npA, and nmA as well as the supplementary data for the case studies are available from github.com/AmiryousefiLab/nmANOVA.

## Supplementary

### S1. Formulating the *F*_*nm*_

For any *α* and *β* in range of *N* number of objects, define *δ*_*αβ*_ as the outcome of the dissimilarity function *δ*(**y**_*α*_, **y**_*β*_) where **y** is either a scalar or a vector of the interested response. Collecting all the pairwise dissimilarities in a square matrix Δ _*N*×*N*_ = {*δ*_*αβ*_} forms the *dissimilarity* matrix between object as the counterpart of *distance* matrix in npA (S3.2). For each *j*′ = 1,2,…,*g* and *j* ≠ *j*′, let’s denote 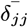 and 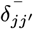 as the mean dissimilarity measures of the diagonal and off-diagonal sub-matrices of the Δ and 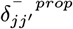 as the mean of a portion of dissimilarity measures in all diagonal matrices, which comprises the same fraction of diagonal matrices as the number of elements in the current sub-matrix in all off-diagonal sub-matrices. It is important to note that 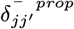 represents the means of proportional fractions that contain elements which do not overlap for any combination of *j* and *j*′. The latter is needed to ensure independence of the sum elements in equation 2. Under the null hypothesis of similarity of all partition means, the *F*_*nm*_ as presented below is following the *F* -distribution with *g*^2^ − *g* and *g* − 1 degrees of freedom;

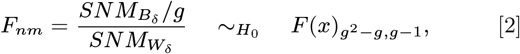

where 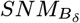 and 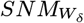 are the *sums of squared normalized means* of dissimilarity values *between* and *within* partitioning groups defined as;

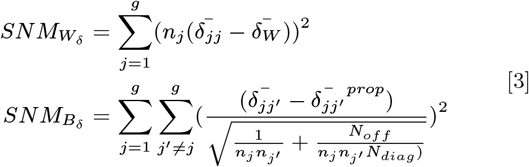

where 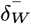 is the overall mean values of all the dissimilarity values in the partitioning groups 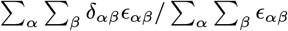, *ϵ* is an indicator function defined in 13, *N*_*of f*_ is the number of elements in all off-diagonal sub-matrices and *N*_*diag*_ is the number of elements in all diagonal sub-matrices.

### S2. Proof of *F*_*nm*_ distribution

Based on the central limit theorem the mean of 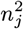 unknown independently distributed *δ*_*αβ*_ with population mean *μ* and variance *σ*^2^ is normally distributed hence,

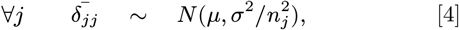

Normalization of these values lead to,

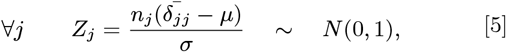

Now under null hypothesis of the sameness of the competing groups mean with each other *H*_0_ : ∀*j,j*′ ∈ (1,, *g*), *μ*_*jj*′_ = *μ* and hence 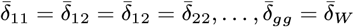, the sum of squared values of the above would be distributed as the *χ*_(*g*−1)_ as,

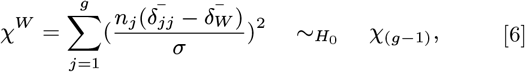

This is true since the sums of squared *g* standard normal random variables is Chi-squared distributed.

Following the same logic for dissimilarities between groups we have,

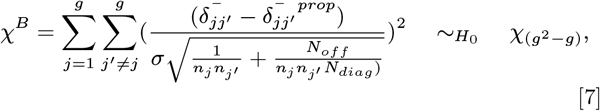

where in that tacitly we have used the equation of the mean values of the dissimilarities *between* groups to be equal with that one as the *within* groups, and for the same reason albeit unlike *χ*^*W*^, we are not losing an extra degree of freedom in *χ*^*B*^. Again with noticing that the division of two Chi-squared distributed divided by their corresponding degrees of freedom is *F* -distributed we have

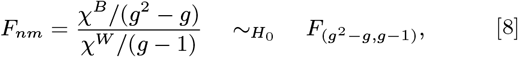

or with rearranging of the values in the above we have,

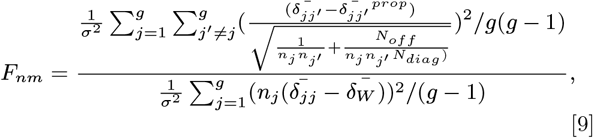

that with crossing out the 1*/σ*^2^ and *g‒*1 the (2) is obtained. Note that the *F*_*nm*_ is capturing the amount of noncentrality incurred by degree of deflection of the *between* mean dissimilarities in contrast to the *within*. To the extend of this deflection, *F*_*nm*_ would be proportionally inflated and if such observed value *F*_*o*_ is more than certain degree 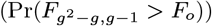, it stands as the existence of evidences for significant differences between the competing groups.

### S3. Essential notes on metric ANOVA

Consider *N* data points, realizations of a random variable Y, and indexed by *α, β, θ* ∈ {1 … *N*}. We further define a metric function *d*, 𝒮 = (*S, d*) as a set of all metric spaces spanned by a metric *d*(*S, S*) : *S* × *S* ↦ [0, ∞) on its domain set *S*. Note that the functions in are not necessarily surjective. The metric function defines distances between data points, *d*(*y*_*α*_, *y*_*β*_). An *N* × *N* distance matrix *D* expresses all pairwise distances between data points. The distances in matrix *D* follow a metric if and only if for all *α, β, θ* ∈ {1 … *N*} the triplets (*α, β, θ*) fulfill the triangle inequality *d*(*y*_*β*_, *y*_*θ*_) ≤ *d*(*y*_*α*_, *y*_*β*_) + *d*(*y*_*α*_, *y*_*θ*_) (18) From swapping the arguments two more equality arise that the collectively with the triangle inequality are defined as metric conditions as:

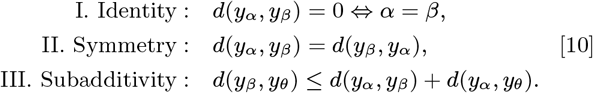

#### 1. Classic ANOVA (cA)

Consider a data set partitioned into *g* groups, each containing *n* _*j❘j* = 1, *…, g*_ items such that *N* = ∑_*j*_ *n*_*j*_ constitute the total number of items. Each item is assumed to be the outcome of a *p*-dimensional random variable, *Y*. For the *k*th dimension, let *y*_*ijk*_ denote the measured response of *i*th object in the *j*th partition, for all *i* = 1, *…, n*_*j*_, *j* = 1, *…, g*, and *k* = 1, *…, p*. The cA tests for the meaningfulness of the partitioning in terms of the proportion of variance explained by using the *decomposition of sums of squares* which partitions the *total* sums of squares *SS*_*T*_ to *between* and *within* groups sums of squares, *SS*_*B*_ and *SS*_*W*_, respectively (7):

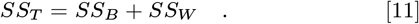

In cA the distances are Euclidean. For the sake of simplicity and without loss of generality, we only use one-dimensional response variable in the following, that is, *p* = 1. In this setup the elements of Equation (11) are defined as:

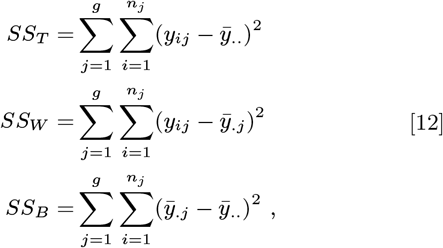

where 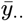 and 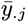 are overall mean of the responses and mean of responses in the *j*th group, respectively.

#### 2. Nonparametric ANOVA (npA)

npA considers pairwise distance matrices between objects (9). The distance matrix is given by a metric function *d* on the responses such that *d* : **Y**×**Y** 1↦ [0, ∞), where **Y** is a realization of a *p* dimensional random variable, given by **y** = {**y**_*ij*_; **y**_*ij*_ = (*y*_*ij*1_, *…, y*_*ijp*_)} ∈ **Y** for all *i* and *j*. We are interested in distances that besides their domain, form an element of 𝒮 ∋ (*Y, d*). All the outcomes of the distance metric function can be collected into a pairwise distance matrix *D*_*N* × *N*_ = {*d*(*y*_*α*_, *y*_*β*_)}, where each entry is the outcome of the distance function for *y*_*α*_ and *y*_*β*_. The applicability of these two methods depends on their different initial assumptions and the available information from the experiment; assuming the same independent variable, cA analyzes response values themselves while the npA analyzes pairwise distances. The latter is more suitable for cases where defining an absolute value for the responses is much more difficult than defining their relative order.

npA results by using the fundamental relationship underlying the topology of a set of points. Specifically, the sum of squared distances between points and their centroid is equal to the sum of squared inter-point distances divided by the number of points. For the set of *N* elements this means that 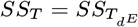, where 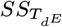 is the mean of the sum of squared Euclidean distances. Analogous to cA, it is possible to partition the sum to *between* and *within* sums of squared distances, 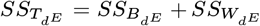. The values of the latter formula are equal to their counterparts in (11). The result generalizes to any general metric function *d*, resulting in the decomposition formula (11)

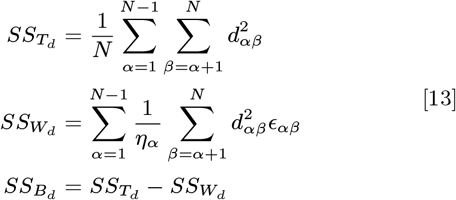

where *η*_*α*_ takes the *n*_*j*_ value if the *α*th observation is in the *j*th group and *ϵ*_*αβ*_ is equal with 1 if both *α* and *β* observations are in the same group and 0 otherwise. Similarly 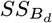 and 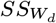 denote the *between* and *within* means of sums of square distances. Note that deriving the 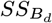 is not straightforward but one can simply make use of the decomposition rule of sums of squares and calculate it as above.

#### 3. Hypothesis testing

In cA, significance of a given partitioning is tested by comparing the averaged ratio of the variance *between* to *within*, hence the *F* -statistic:

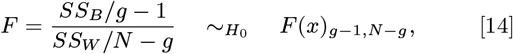

which under null hypothesis follows *F* (*x*)_*g*−1,*N* −*g*_ distribution. We refer to the values of test statistic based on the observed outcomes of the experiments as *F*_*o*_. This value then will be compared against the null *F* distribution to obtain the *P* -value, *P* = Pr(*F*_*g*−1,*N* −*g*_ *> F*_*o*_). Rejection of the null hypothesis indicates the presence of at least one group with mean value differing from the others. With the similar indication for the p-value, in npA a pseudo permutative *F* -statistic is defined as

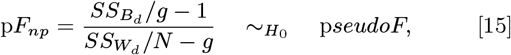

Indexing each values of p*F*_*np*_ obtained from *π* permutation as p 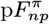 the *P* -value can be obtained as *P* –value 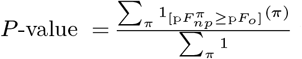 where 1_[*A*]_(*x*) is an indicator function (that is 1 if *x* ∈ *A* and 0 otherwise), and p*F*_*o*_ is the observed pseudo *F* -statistic.

#### S4. Data

The R code for implementing the nmA algorithm as well as the supplementary data SD1, SD2, and SD3 are available from https://github.com/AmiryousefiLab/nmANOVA.

